# Prior exposure to pathogens augments host heterogeneity in susceptibility and has key epidemiological consequences

**DOI:** 10.1101/2024.03.05.583455

**Authors:** Dana M. Hawley, Anna A. Pérez-Umphrey, James S. Adelman, Arietta E. Fleming-Davies, Jesse Garrett-Larsen, Steven J. Geary, Lauren M. Childs, Kate E. Langwig

**Affiliations:** Department of Biological Sciences, Virginia Tech, Blacksburg, VA, USA; Department of Biological Sciences, University of Memphis, Memphis, TN, USA; Department of Biology, University of San Diego, San Diego, CA, USA; Department of Pathobiology & Veterinary Science, University of Connecticut, Storrs, CT, USA; Department of Mathematics and Virginia Tech Center for Mathematics of Biosystems, Virginia Tech, Blacksburg, VA, USA

## Abstract

Pathogen epidemics are key threats to human and wildlife health. Across systems, host protection from pathogens following initial exposure is often incomplete, resulting in recurrent epidemics through partially-immune hosts. Variation in population-level protection has important consequences for epidemic dynamics, but how acquired protection influences inter­individual heterogeneity in susceptibility and its epidemiological consequences remains understudied. We experimentally investigated whether prior exposure (none, low-dose, or high­dose) to a bacterial pathogen alters host heterogeneity in susceptibility among songbirds. Hosts with no prior pathogen exposure had little variation in protection, but heterogeneity in susceptibility was significantly augmented by prior pathogen exposure, with the highest variability detected in hosts given high-dose prior exposure. An epidemiological model parameterized with experimental data found that heterogeneity in susceptibility from prior exposure more than halved epidemic sizes compared with a homogeneous population with identical mean protection. However, because infection-induced mortality was also greatly reduced in hosts with prior pathogen exposure, reductions in epidemic size were smaller than expected in hosts with prior exposure. These results highlight the importance of variable protection from prior exposure and/or vaccination in driving population-level heterogeneity and epidemiological dynamics.

## Introduction

Pathogen epidemics are increasing in frequency in humans and other animals (1,2), underscoring the need to characterize key sources of heterogeneity that influence epidemiological dynamics for pathogens. Hosts in a population often show considerable heterogeneity in contact rates (e.g. 3,4), infectiousness (e.g. 5), and even pathogen susceptibility (e.g. 6), defined here as the probability of infection given exposure. The individual­level traits that predict host susceptibility to a given pathogen, including sex (7), genotypic variants (8), and immune history (9), have been widely described for some systems. Nonetheless, the coarse discretization of these traits into mathematical models often inaccurately describes the extent of heterogeneous infection risk present among hosts in a population (10–13). A key approach to quantifying such heterogeneous risk at the population level is to estimate the shape of the distribution of host susceptibility (14–16), either using epidemiological data (e.g. 17,18) or experimental studies of non-human animals (e.g. 19,20). Overall, such population-level approaches to quantifying host susceptibility distributions are particularly important because the shape of such distributions, and particularly the extent of inter-individual heterogeneity present in a host population, can have key downstream consequences for epidemic dynamics (e.g. 14,16,21,22), Thus, it is critical to understand how factors such as acquired protection from prior pathogen exposure alter the degree of population­level heterogeneity in susceptibility, and its epidemiological consequences.

Host reinfection is a common but understudied feature of many host-pathogen systems (23–28), including SARS-Cov-2 (29), pneumococcal disease (30), influenza (31), malaria (32), and other diseases of agricultural and wildlife significance. Because the protection acquired from prior pathogen infection is often incomplete and/or wanes over time, reinfections occur even in systems where hosts have lower mean susceptibility during secondary exposures, relative to individuals exposed for the first time (33–36). While there is growing appreciation for the pervasiveness of heterogeneity in acquired host protection in response to vaccination or infection across systems (31,35,37,38), prior work has largely focused on how host protection from vaccination or prior infection influences mean population traits, rather than variability among individuals in a given population (10). As such, it remains unknown how host protection acquired from prior pathogen infection alters the shape of the susceptibility distribution present in a given host population.

Prior pathogen exposure can modify inter-individual heterogeneity in susceptibility via several mechanisms. First, prior exposure to pathogens could generate incomplete (or “leaky”) protection similarly for all exposed hosts (35), such that all exposed hosts acquire identical levels of protection against reinfection. Alternatively, prior pathogen exposure could result in relatively complete acquired protection for some proportion of hosts, while others remain relatively unprotected (termed “all-or-nothing” immunity (35)). Epidemiological models often select either wholly leaky protection of hosts that allow every host an identical rate of reinfection, or all-or-nothing protection such that some hosts remain in the recovered class while a subset is routed back to the susceptible class, representing two extremes of how heterogeneity in susceptibility is incorporated into disease models (10,20). Whether and how prior exposure to a pathogen alters the shape of the susceptibility distribution for a given host population likely depends on the mode of action of acquired protection [“leaky”, “all-or-nothing” (10,20), or some intermediate (38)], as well as the extent of prior pathogen exposure that hosts experience (39). For example, host exposure to low pathogen doses, which can occur during the natural transmission process for many pathogens (e.g. 40-42), might be more likely than higher exposure doses to generate incomplete host protection (43,44) and thus to induce population­level heterogeneity in susceptibility.

Regardless of the underlying mechanisms, effects of prior exposure on population-level heterogeneity in host susceptibility are particularly important to characterize because this heterogeneity can have key epidemiological consequences, including lower predicted outbreak sizes (20) and higher rates of reinfection (22). This is in part due to the process of cohort selection, through which the most susceptible individuals are infected first in any given epidemic time step, leaving lower mean susceptibility among remaining uninfected hosts in a population (45). Experimental quantification of population-level heterogeneity in susceptibility draws on quantitative microbial risk assessment approaches (46) to estimate susceptibility distributions using a series of increasing pathogen challenge doses. Because only the most susceptible hosts become infected at low challenge doses, whereas even the least susceptible hosts become infected at the highest challenge doses, dose-response models can be used to estimate the degree of host heterogeneity in susceptibility in a given population (20). Importantly, by using controlled pathogen exposure doses, experimental studies allow direct quantification of host heterogeneity in susceptibility without the confounds of variation in exposure, a key additional source of heterogeneity in free-living populations (47). Thus, dose­response studies of non-human animals are particularly useful for isolating effects of factors such as vaccination (20) or maternal parasite exposure (19) on the degree of heterogeneity present in a host population as a whole. Here we apply such experimental approaches to ask how prior pathogen exposure alters the degree of inter-individual heterogeneity in susceptibility. Robustly quantifying changes in population susceptibility distributions that occur with prior pathogen exposure is challenging due to large sample size requirements of experimental dose response studies. Nonetheless, studies of host susceptibility distributions are critical for adequate characterization of host-pathogen dynamics, including the potential downstream effects of inter-individual heterogeneity on pathogen strain coexistence and virulence evolution (21,48,49).

Here we investigate effects of host prior exposure on inter-individual heterogeneity in susceptibility using a tractable wildlife-pathogen system: a common North American songbird species (house finch; *Haemorhous mexicanus*) and its bacterial pathogen, *Mycoplasma gallisepticum* (MG), which causes severe conjunctivitis. The potential effects of prior pathogen exposure on population-level heterogeneity in susceptibility are particularly relevant in this system because MG exposure doses are variable across free-living hosts: susceptible birds contact MG that was deposited onto bird feeder surfaces in variable amounts by infected individuals (50,51). Further, although the conjunctivitis caused by MG indirectly results in significant mortality in finches, free-living hosts recover at high rates from initial infection (52), leaving a pool of recovered hosts in natural populations. Experimental studies show that recovered hosts have significant acquired protection relative to pathogen-naive hosts, but can still be successfully reinfected at high rates, even with homologous pathogen strains (39,53). Finally, our work to date suggests that experimental variation in the degree of prior exposure to MG (in both dose and number of exposures) results in heterogeneous protection from reinfection among finches, as measured by conjunctival pathogen loads following reinfection challenge (39,43,44). Whether prior exposure to MG specifically alters inter-individual heterogeneity in susceptibility, a population-level trait that requires dose-response approaches to robustly quantify, has not yet been examined.

To test how variation in prior pathogen exposure alters population-level distributions of susceptibility, we first experimentally varied the degree of prior pathogen exposure categorically (none, low-dose, or high-dose) in wild caught-finches that we confirmed had no exposure to MG prior to capture. After recovery from prior exposure treatments, we measured host susceptibility (0|1) to secondary dose challenge (Figure 1), fitting dose-response models for each treatment group to determine whether host prior exposure treatment altered the degree of population-level heterogeneity in susceptibility. Specifically, for each treatment group, we considered population­level distributions of susceptibility that represent either identical infection probabilities (homogeneous model) or models that represent inter-host variation in rates of infection (heterogeneous model, where numerical susceptibility values were estimated according to a fitted gamma distribution (16,19,20,54)). Importantly, our models allowed us to disentangle effects of prior exposure on mean susceptibility from changes in heterogeneity in susceptibility. Lastly, to determine effects of exposure-induced population heterogeneity on resulting epidemic dynamics, we parameterized an SIR model for this system with susceptibility distributions and expected mortality rates in the wild estimated from our experimental data.

**Figure 1.**
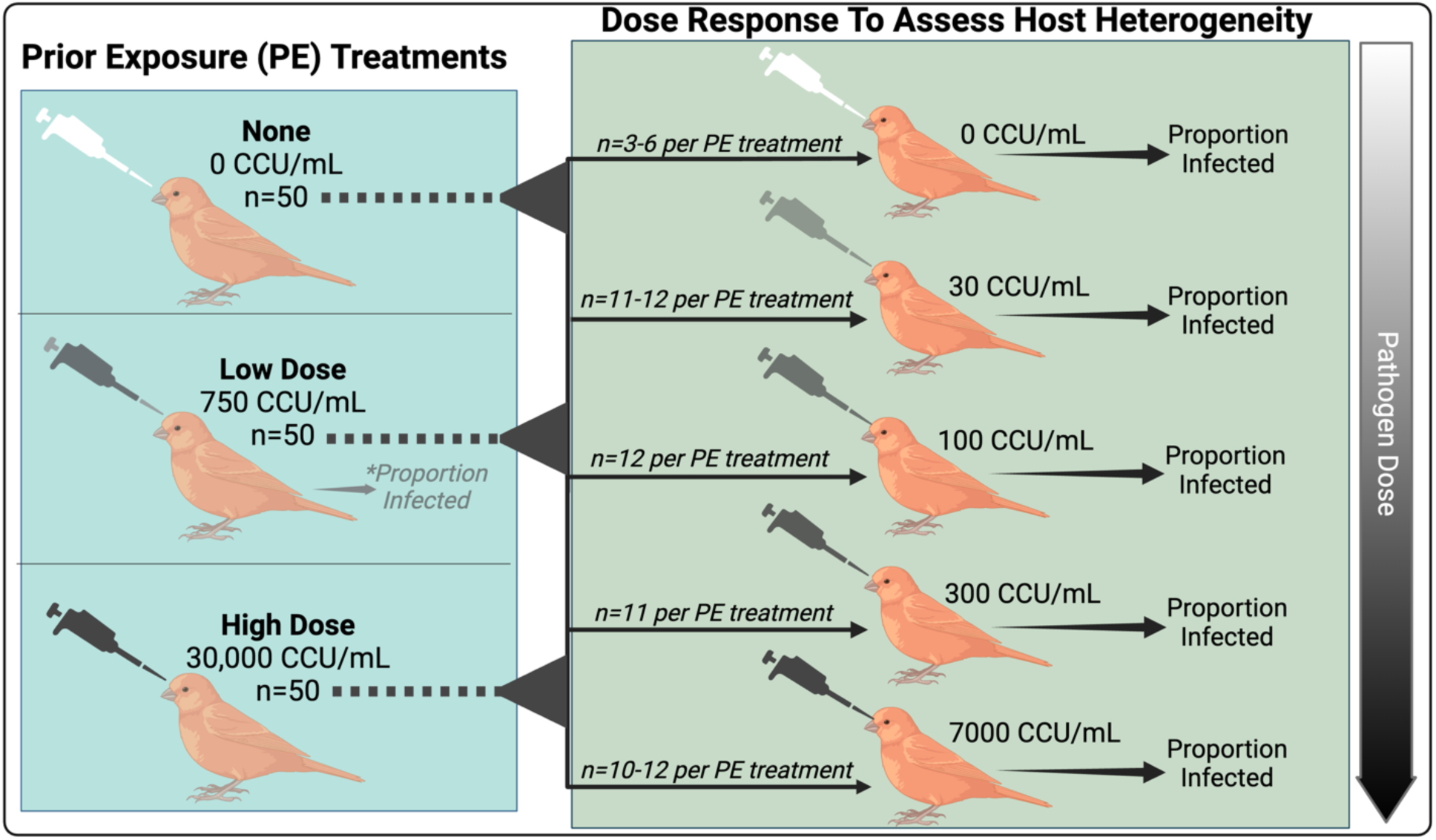
Experimental design for quantifying how prior exposure to *Mycopiasma ga/lisepticum* alters host heterogeneity in susceptibility in house finches (n=150 for final analysis). Five weeks after one of three prior exposure (PE) treatments (none, low, or high dose; left panel), all birds received a secondary dose challenge (right panel) to assess heterogeneity in susceptibility. Our primary data set was whether birds became infected (0|1) in response to a given secondary dose (right). ‘However, to improve model fits, we also used bird responses to low-dose PE treatment (left, asterisk), which fell intermediate to our highest secondary challenge doses (300 and 7000 Color Changing Units [CCU]/mL), to quantify the proportion of birds with no prior exposure (at the time of low-dose PE) that become infected at a 750 CCU/mL dose. Made in Biorender.

## Results

### Prior exposure to pathogens augments population-level heterogeneity in susceptibility

Birds with no prior pathogen exposure at the time of secondary dose challenge had low variability in host susceptibility (coef of variation [CoV] from gamma distribution = 0.899) and there was no support that heterogeneous dose-response models were an improvement over models assuming homogeneous host susceptibility (defined here as identical infection probabilities across hosts in a population) (Fig 2a; deviance homogeneous = 1.248; deviance heterogeneous = 3.307). Birds with no prior pathogen exposure also had high mean susceptibility to infection (mean susceptibility [fitted gamma distribution]=1.181). In contrast, dose-response curves were shallower and fewer birds were infected upon secondary exposure in both the low (mean susceptibility = 0.446 [fitted gamma], CoV [fitted gamma] = 1.630; Fig 2b), and high-dose (mean susceptibility = 0.192 [fitted gamma], CoV [fitted gamma] = 2.511; Fig 2c) prior exposure groups. Both prior exposure treatments were better described by models assuming heterogeneous versus homogeneous susceptibility (Fig 2; likelihood ratio tests heterogeneous versus homogeneous (46), low dose: P = 0.020, high dose: P = 0.033).

**Figure 2.**
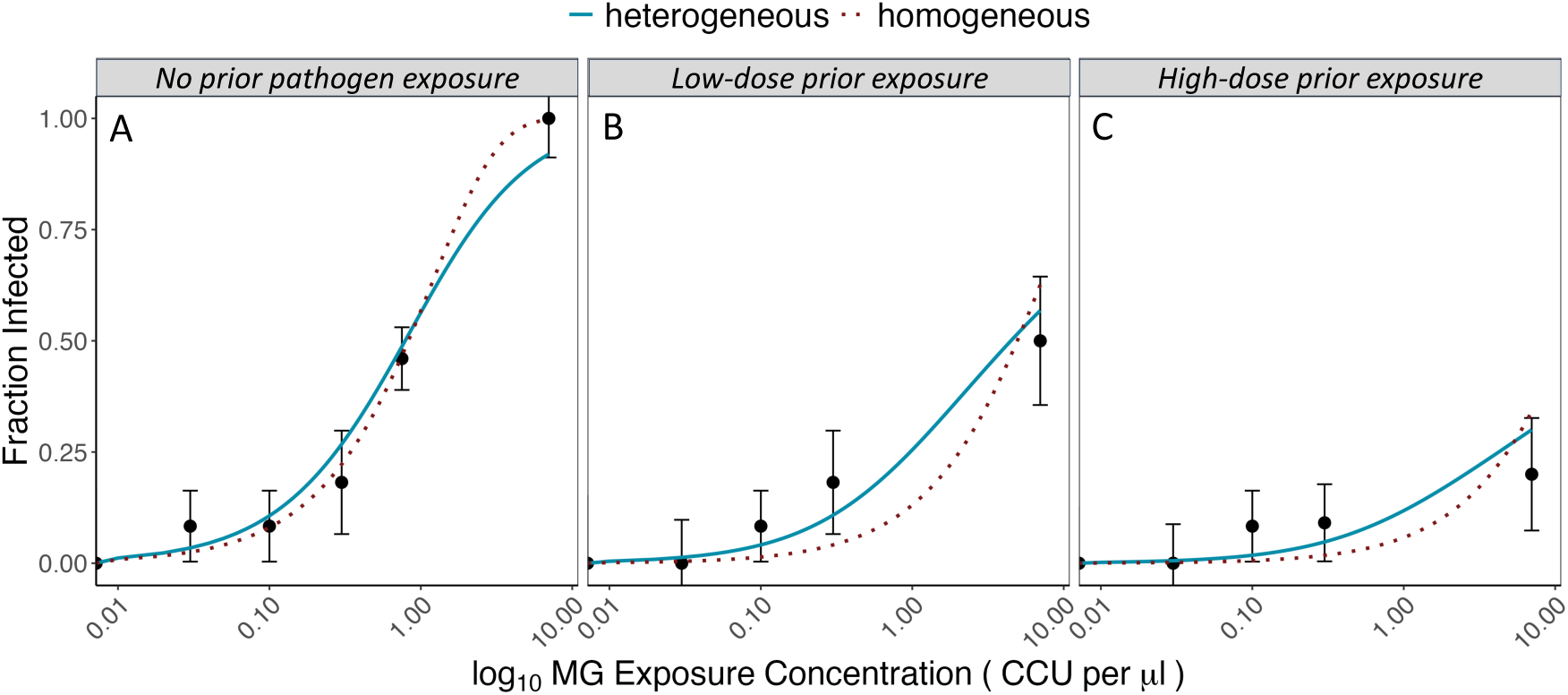
Dose response curves for house finch susceptibility to secondary challenge with *Mycoplasma gallisepticum* across prior exposure treatments. Points (+/− 1SE) show the fraction of individual birds (n=10-12 birds for most points; individual responses are 0|1) infected at each secondary exposure dose, shown as Color Changing Units (CCU)/uL. Lines indicate model fits, with blue indicating gamma (heterogeneous) model fits, and red dashed lines indicating homogeneous model fits. Panel labels show prior exposure treatment (birds in the no prior exposure treatment were pathogen-naive at the time of secondary dose challenge). In hosts with prior pathogen exposure (low and high-dose prior exposure groups), the gamma model (which accounts for inter-individual heterogeneity) was better supported via likelihood ratio tests.

To estimate effects of prior exposure on host mortality rates in the wild, we used clinical scores of eye disease in response to secondary dose challenge to estimate mean mortality rates for each prior exposure treatment group (as per (34); see *Methods*). As expected, predicted host mortality rates decreased with the degree of prior host exposure (Table S2), declining from 0.05894/day in birds with no prior exposure, to 0.02806/day in birds with prior low-dose exposure, and to a negligible value (2.483*10^-17^) in birds with prior high-dose exposure.

### Exposure-induced heterogeneity in susceptibility reduces epidemic size

Using these empirically-derived susceptibility distributions and predicted mortality rates (Fig 3, Table S2) to parameterize an SIR model of this system, we find that the total epidemic size is largest (97.13% of the population) for a host population with no prior pathogen exposure, while for low and high-dose prior exposure groups, the total epidemic size is significantly reduced (33.67% and 16.13% of the population, respectively; blue bars, Fig 4A). Because prior pathogen exposure also protects hosts from mortality, there is no mortality during the epidemic in the high-dose prior exposure population, while there is 16.28% mortality in the low-dose prior exposure population, and more than 64.37% mortality in the naive host population (darker colored portion of bars, Fig 4A; dashed lines, Fig 4C).

**Fig 3.**
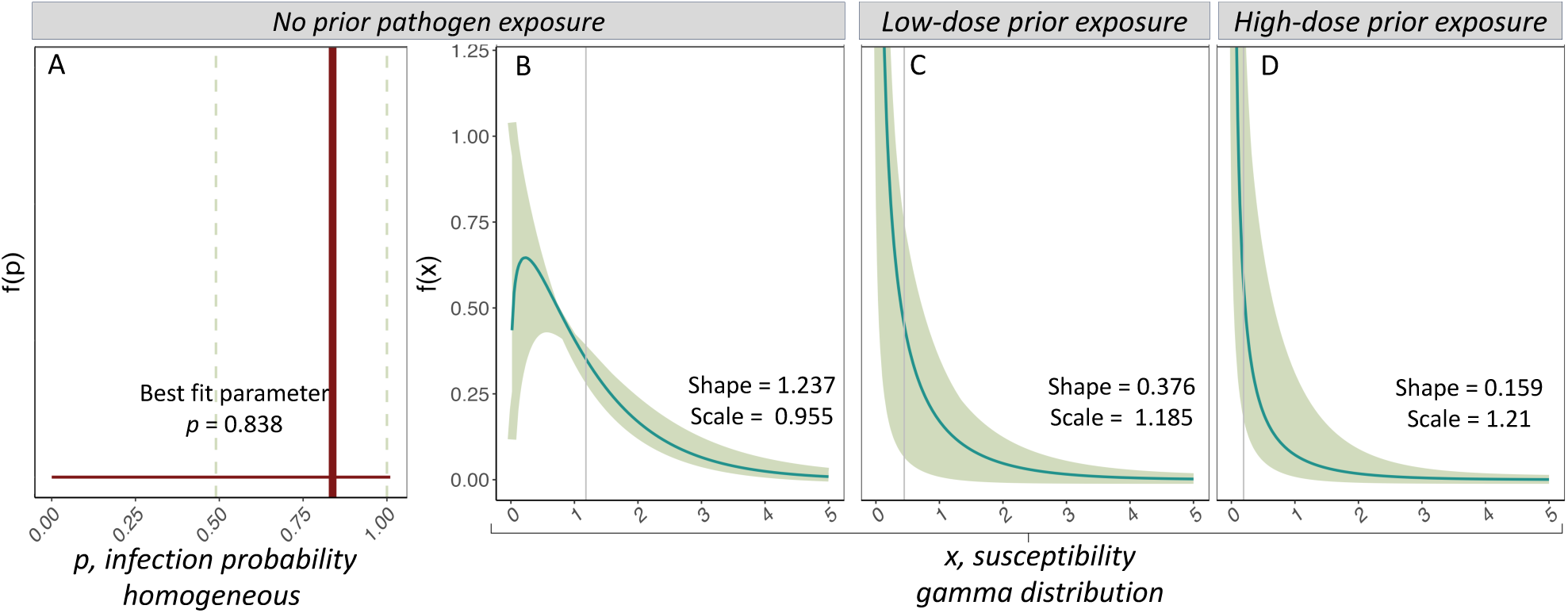
Host susceptibility distributions for house finches from variable prior exposure treatments: no prior exposure (A,B); low-dose (C); or high-dose (D). Colored lines show estimated susceptibility distributions from either homogeneously (A) or gamma-distributed (B-D) models (note distinct axes for the two models). In (A), host infection probability per 1000 bacterial particles (p) is shown as the single best fit parameter *p* (dotted vertical lines represent 1 standard error) for the homogeneous model, which was the best fit model for the no prior exposure group (see *Results).* In (B-D), the best fit parameters (shape and scale) for gamma distributions (teal lines) are listed for each group, and vertical gray lines indicate mean susceptibility (*x*) for that treatment. Lighter shading represents 95% confidence regions for gamma distributions, obtained by bootstrapping chi-squared residuals to create 1,000 pseudoreplicates of infection data and then refitting the model to pseudoreplicates, as per (20,46). The gamma model was the best fit for only the low-dose and high-dose groups. Gamma estimates are also shown for the no prior exposure group (B) because this allowed more equivalent comparisons for certain SIR simulations (see *Methods*).

**Fig 4.**
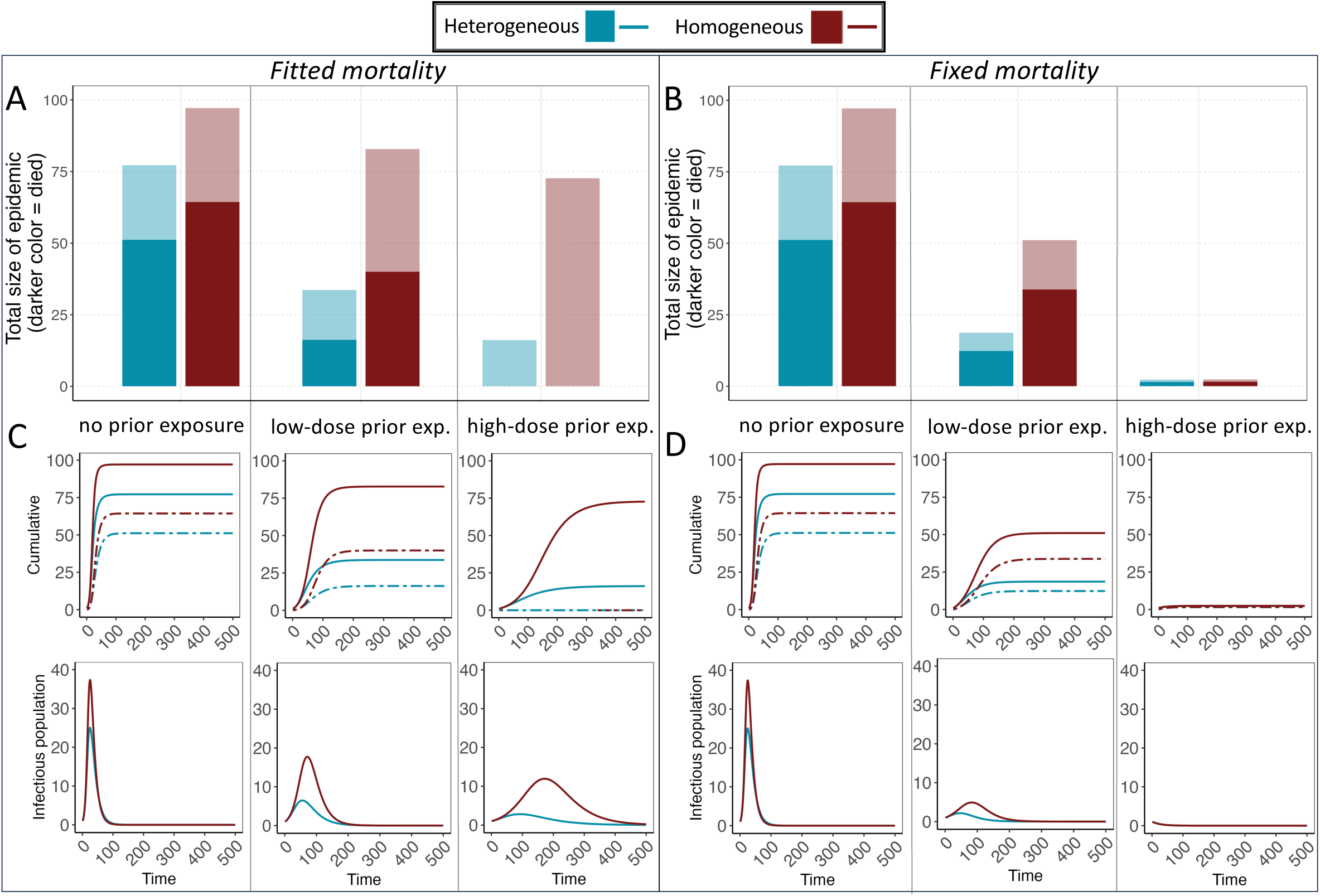
Cumulative epidemic size (proportion of individuals who ever entered the infectious class) (A,B); dynamics of cumulative infections and deaths (top: C, D; deaths: dashed lines) over time, and numbers of infectious hosts over time (bottom: C, D) in our SIR model, as a function of prior exposure (abbreviated as “exp.”) treatment (x-axis for A,B; columns for C,D). Total starting population was 100; darker shading within color represents individuals (within cumulative totals) that did not survive the epidemic. Blue bars and lines use fitted heterogeneous susceptibility (gamma distribution) for all prior exposure groups (Fig 3, Table S2). Red bars and lines assume homogeneous susceptibility for all groups, with mean susceptibility for each group equal to that of the fitted heterogeneous distribution for that treatment group. Left panels (A,C) use fitted mortality rates, whereby estimated mortality rates declined with prior exposure, while right panels (B,D) assume the fixed mortality rate of all groups is equal to that of the no prior exposure birds.

We also examined whether the observed reductions in epidemic size with host prior exposure are driven primarily by changes in the susceptibility distribution, versus reductions in mean host susceptibility (blue versus red bars within each prior exposure treatment group, Fig 4A, B). For empirically determined best fit parameters, epidemic size for a paired simulation assuming homogeneous rather than heterogeneous susceptibility (while holding mean susceptibility constant) was larger by 49% and 56%, under low dose and high dose prior exposure, respectively (low dose, fitted mortality: 49.12 epidemic size difference [hom-het]; high dose, fitted mortality: 56.55 epidemic size difference [hom-het]). We simulated the heterogeneous and homogeneous models using the parameter estimates obtained from bootstrapping chi-squared residuals (see Fig 3). Assuming homogeneous versus heterogeneous susceptibility (while holding mean susceptibility constant for a given prior exposure treatment) results in consistently larger outbreaks (Fig S1; difference in epidemic size = homogeneous [red]-heterogeneous [blue] outbreak size for a given mean susceptibility). Indeed, across all simulations, outbreaks are never smaller in models assuming homogeneous versus heterogeneous susceptibility (all P < 0.0001; Fig S1; epidemic size differences [95% CIs of hom-het epidemic size] for each simulation: low dose, fitted mortality [2.078,49.12]; high dose, fitted mortality [0.8994,61.29]; low dose, fixed mortality [0.1187,40.77]; high dose, fixed mortality [0.010,36.35]). Together, these results indicate effects of host heterogeneity *per se* on outbreak size, which act above and beyond the effects of lower mean susceptibility from prior pathogen exposure.

Reductions in mean susceptibility with host prior exposure also contribute to smaller outbreak sizes, but the extent depends on the assumed level of host mortality. When controlling for mortality differences (i.e., all populations are assumed to have mortality equivalent to the naïve, no-prior exposure population), the outbreak size of both groups with prior pathogen exposure (low or high) is reduced (Fig 4B,D). In the case of simulations assuming homogeneous susceptibility (red bars, Fig 4B) while accounting for differences in mean protection with prior exposure, this reduction is *entirely* due to lower mean host susceptibility resulting from prior pathogen exposure. However, when heterogeneity in susceptibility and reduced mean susceptibility are both accounted for, further reductions in outbreak size and mortality are found for the low-dose prior exposure group (blue bars and lines, Fig 4B,D). Further reductions from heterogeneity *per se* are not possible for the high-dose prior exposure group, for which our model produced an outbreak size of nearly 0 from changes in mean susceptibility alone.

In the more biologically realistic setting (Fig 4A,C) with changes in susceptibility and host mortality in response to prior exposure both accounted for, we find a more complicated pattern. If we assume a homogeneous population (red bars) with the mean susceptibility found empirically for low and high-dose prior exposure groups (and mortality rates set as empirically determined values), the epidemic sizes drastically increase (82.79% and 72.68% of the population for the low and high prior exposure, respectively) relative to models accounting for host heterogeneity in susceptibility (blue bars and lines; Fig 4A,C). The large outbreak in the high-dose prior exposure population (despite very low mean susceptibility in this group) is due to the absence of disease-induced mortality in this population, resulting in a higher basic reproductive number because all individuals survive the entire infectious period.

## Discussion

Here we use a naturally-occurring host-pathogen system characterized by high rates of reinfection (43) to test how acquired protection from variable degrees of prior pathogen exposure influences population-level distributions of host susceptibility. Using experimental infection data for birds across three distinct prior exposure treatments, we tested for population­level heterogeneity by statistically comparing dose response models that assume either identical susceptibilities among hosts in a population (homogeneous model) or inter-host variation in susceptibility (heterogeneous model). We find that prior exposure to either low or high doses of pathogen significantly augments the extent of population-level heterogeneity in susceptibility relative to a pathogen-naive host population, with population-level heterogeneity increasing with the degree of prior pathogen exposure, from none, low, to high-dose. In addition, our SIR model demonstrates that the observed changes in the host susceptibility distribution with prior pathogen exposure have key effects on epidemic dynamics. After accounting for estimated effects of prior exposure on host mortality, changes in population-level heterogeneity in susceptibility had stronger effects on outbreak dynamics than reductions in mean host susceptibility, resulting in dramatically lower outbreak sizes.

The past several decades have brought growing recognition that many host-pathogen systems are characterized by reinfection potential, whether shortly following initial infection and recovery, or over longer timescales after any host acquired protection from prior infection has waned (23–26,29,55–58). Despite the ubiquity of highly specific immune memory that allows hosts to respond rapidly to and effectively resist reinfection with the same pathogen, the acquired protection generated by prior pathogen exposure is often incomplete, even in vertebrate taxa with highly specific antigen repertoires and associated immune memory (59). Thus, some degree of reinfection is still possible, despite lower mean susceptibility in hosts with acquired protection (33,48). Given the importance of reinfection in many systems, it is critical to characterize variability in key epidemiological traits for host populations made up of individuals with some degree of prior pathogen exposure. For example, two longitudinal studies of repeated malarial episodes in children documented substantial heterogeneity in the length of time until *Plasmodium falciparum* reinfection (60) among children and the degree of acquired protection generated against clinical malaria (60,61). However, this heterogeneity was attributed primarily to variation among children in the extent of repeated exposure to the parasite. Because such heterogeneity in exposure to pathogens is common in natural populations (47), it can be challenging to quantify effects of acquired protection on heterogeneity in host susceptibility *per se* without the use of controlled experiments. Nonetheless, it is critical to characterize effects of acquired protection on host susceptibility distributions in particular, because population-level heterogeneity in susceptibility can influence the likelihood and severity of outbreaks (3,10,15,62), as well as the evolution of virulence for pathogens (21,48,49,63).

One intuitive prediction is that host acquired protection from prior exposure to pathogens should homogenize variation in susceptibility in a population by reducing susceptibility to reinfection similarly for all individuals. Instead, we found that the protection acquired from prior exposure to a naturally-occurring bacterial pathogen, whether at low or high doses, significantly augmented population-level heterogeneity in susceptibility in house finches. Importantly, while our dose-response estimates for birds with no prior exposure to MG are well supported by models assuming homogeneous susceptibility, this does not indicate that there is no biological variation in susceptibility present within this treatment group. Instead, this result suggests that variation in susceptibility among house finches without prior pathogen exposure is of low magnitude, which is further supported by the low coefficient of variation of the estimated gamma distribution for this group. Prior studies in this system find notable variation among both individuals and populations in other types of host responses to experimental MG infection, including infection severity (pathogen loads), disease severity (eye scores), and tolerance (per­pathogen disease severity) (e.g. 64-67), even for birds with no prior exposure to MG infection. Whether host susceptibility *per se* tends to be less variable than other types of host responses, both in this system and others, is an interesting topic for future inquiry, particularly given the key epidemiological consequences of heterogeneity in host susceptibility (14–16).

Few studies have explicitly tested how population-level distributions of host susceptibility change in the presence of prior pathogen exposure or host protection. However, two experimental studies that examined how other forms of acquired protection (transgenerational pathogen exposure, and vaccination) influence the shape of host susceptibility distributions found surprisingly similar results to what we report here. First, Ben-Ami et al. (19) showed that maternal exposure to a high dose of a bacterial pathogen in *Daphnia magna* widened the susceptibility distribution of the offspring without affecting its mean. Second, Langwig et al. (20) found that vaccination of rainbow trout against a virus augmented population-level heterogeneity in susceptibility to that virus relative to unvaccinated individuals, while also reducing mean susceptibility. Interestingly, the results from this vaccination study qualitatively match what we found for house finches, even though the protection that hosts acquire from active infection can be stronger and/or more variable in strength or duration than that acquired by vaccination (e.g. 68,69). Further, our detected pattern of increasing heterogeneity in susceptibility with higher degrees of prior pathogen exposure, from no, low-dose, to high-dose prior exposure treatments, suggests that stronger stimulation of host acquired immune responses by high pathogen doses (in this case, 30000 CCU/mL) can, at least in this system, lead to even more variability across individuals in a population. Intriguingly, we observed highly variable responses to prior exposure despite using an MG strain (VA1994) isolated from free-living house finches over 25 years ago. Because significant host evolution has occurred since then (64,67), one might expect less variable host responses to the strain used here, compared to a more recent MG strain, if host resistance traits have been favored or even fixed in the host population (70). Nonetheless, prior exposure to an older, basal house finch MG strain still induced significantly higher levels of host heterogeneity in susceptibility, even at low prior exposure doses. Because strain traits such as virulence may influence the degree of host heterogeneity in a population (48), future work should examine whether prior exposure to MG strains more virulent and derived than the strain used here generates even higher degrees of heterogeneity in susceptibility in a host population.

We used an SIR model parameterized empirically to predict how the population-level susceptibility distributions induced by host prior exposure would influence epidemic dynamics in this system. Consistent with prior work using SIR (20,22) and dynamic network approaches (71), we show that population-level heterogeneity in susceptibility (here as a result of prior pathogen exposure) suppresses overall outbreak size relative to models that assume homogeneous host susceptibility. Importantly, under either set of mortality assumptions used in our model, outbreak size is always reduced when prior exposure generates heterogeneity in susceptibility, compared to the homogeneous case that still accounts for reductions in mean host susceptibility with prior exposure. These results indicate direct effects of exposure-induced heterogeneity *per se* on outbreak size, effects which act above and beyond effects of mean susceptibility from prior pathogen exposure on outbreak size. Moreover, under the more realistic assumption of host protection from mortality after prior pathogen exposure, the reductions on overall epidemic size in our models appear to be driven predominantly by increased population­level heterogeneity in susceptibility. This observation is consistent with herd immunity derivations in a similar SEIR model, which finds strong dependence on the coefficient of variation (72).

An earlier SIR model of vaccine-induced heterogeneity similarly found that augmented population-level heterogeneity in susceptibility reduced outbreak size (20), but this model did not incorporate potential changes in host mortality rate. Reductions in mortality rate in hosts with prior pathogen exposure, which we estimated were significant in the house finch system, are likely common in hosts with acquired protection, because such protection often reduces host disease and mortality more strongly than infection risk (e.g. 73,74). In our model results, the degree to which prior exposure-induced reductions in mean susceptibility drive outbreak size depends on assumptions of mortality rates in our model. In our model with empirically-derived mortality rates, infected birds in the prior exposure groups are more likely to survive infection (lighter color sections of bars; Fig 4A). Intriguingly, these higher survival rates nearly outweigh the protective effects of reduced mean susceptibility in previously-exposed hosts, resulting in relatively little reduction in epidemic size in both prior exposure groups (assuming homogeneous susceptibility for both groups; red bars, Fig 4A), despite significantly lower mean host susceptibility in birds with prior exposure. In contrast, when mortality rates are held constant regardless of prior exposure treatment, exposure-induced reductions in mean host susceptibility alone result in substantially smaller outbreak sizes (red bars; Fig 4B). These results indicate that effects of prior exposure on mortality rate and susceptibility distributions are both critical to account for in epidemiological models, as they may interact to drive outbreak size.

One limitation of our SIR model is that it assumes that epidemics begin in host populations entirely composed of individuals of one prior exposure type. While such models are likely realistic representations of host populations at certain times of the year (e.g., epidemics in late summer that occur in juvenile flocks composed of fully susceptible finches (75,76)), future models should also consider scenarios whereby populations are composed of some individuals with no prior exposure, and others with varying degrees of prior pathogen exposure. Developing a simulated population to such a state would require knowing how the level of susceptibility of individual birds is altered by prior exposure, for which no data exists. Another limitation is the choice of an SIR rather than an SIRS model that allows for loss of immunity over time, as is the case in the house finch-MG model system, in which protection wanes over approximately one year (53). To disentangle effects of population-level heterogeneity of susceptibility induced by prior exposure, we focus this study on a single, short timescale epidemic while ignoring demographic effects and re-infection, similar to previous work (38,77,78). Allowing for reinfection in our model would require an understanding of how susceptibility and infectivity are linked, because such correlations have an impact on the progression of epidemics (77,78). However, such correlations are particularly challenging to quantify in natural host-pathogen systems. Overall, our work represents an important first step in incorporating one key source (prior exposure) of population-level heterogeneity in susceptibility into mathematical models. Although the importance of individual variation in infection-derived immunity for epidemiological dynamics is increasingly recognized (31,35,79), to our knowledge, our study is the first to explicitly quantify how acquired protection from prior pathogen infection influences population­level distributions of host susceptibility.

In summary, our results highlight the key importance of variable protection from prior pathogen exposure in driving host heterogeneity in susceptibility, a population-level trait with key downstream consequences for epidemiological and evolutionary dynamics (15,22,48). Because the host protection acquired from a prior infection (or vaccination) is often incomplete and/or wanes over time in diverse vertebrate systems (e.g. 29,58), infection-induced heterogeneity in susceptibility is likely a common phenomenon for many animal populations, including humans. While work to date has largely focused on characterizing innate, individual­level sources of host heterogeneity in susceptibility [sex (80), genotype (81), etc.], our results show that population-level heterogeneity in susceptibility can be readily “induced” by prior pathogen exposure, with key downstream consequences.

## Methods

### Bird Capture and Housing

We captured 157 mixed-sex, hatch-year house finches (aged by plumage (82)) in Montgomery County, Virginia using a combination of cage traps and mist nets in June-Aug 2021. Capture and collection were approved by Virginia Department of Game and Inland Fisheries (066646) and USFWS (MB158404), and all handling and care procedures were approved by the Virginia Tech (VT) IACUC. Birds were housed in pairs for a two-week minimum quarantine period in an indoor animal facility, and captured every 3-4 days to assess potential clinical signs of mycoplasmal conjunctivitis. Any birds with signs of conjunctivitis (see below) were immediately isolated, along with their cage-mates, and were not used in experiments.

For all birds that never showed clinical signs of conjunctivitis during quarantine, blood samples were collected 14-18 days post-capture and run for ELISA via a commercial IDEXX kit (99–06729) using methods previously described (83). Only birds seronegative via ELISA on days 14-18 post-capture were assigned to MG-exposed treatment groups (Table S1; all birds except n=3 “prior sham-0 challenge” birds; see Supplement). All birds were housed individually and then resampled one week before prior exposure treatments for final confirmation of seronegativity just prior to experiment initiation.

### Experimental Design and Timeline

We created variation in prior exposure using three categorical treatments (Figure 1, Table S1): no prior exposure to MG (inoculation with the same volume of sterile Frey’s media as a “sham” treatment), low-dose prior exposure (dose of 750 Color Changing Units [CCU]/mL of MG), or high-dose prior exposure (dose of 30,000 CCU/mL of MG). Birds were infected with an MG strain (VA1994 isolate of MG; [7994-1 (6P) 9/17/2018]) that was isolated shortly following the pathogen’s emergence in house finches; despite significant host and pathogen evolution since then (64,84,85), this 1994 strain still results in pathogen load and eye scores trajectories that are qualitatively similar to later-isolated strains (VA2013 (64)). Birds were inoculated with 70 *uL* of their assigned treatment concentration directly into the conjunctivae via micropipette droplets, and then allowed to recover (with recovery quantified at day 41 post-infection; see below).

Forty-two days after prior exposure treatment, birds were given a secondary dose challenge with the same MG strain (again, 70uL directly inoculated into the conjunctivae). Using a fully factorial design (Table S1), we inoculated birds from each prior exposure treatment with one of five MG exposure doses to assess heterogeneity in susceptibility (Figure 1). We selected prior exposure treatments and challenge doses (also termed “secondary doses”) most likely to generate and detect differences in susceptibility based on our published work on conjunctival loads (44), and we minimized other inherent sources of variation in susceptibility in our wild-caught birds by using same-age birds, excluding birds with detectable prior exposure at capture, and randomizing assignments of sex within treatment groups, such that sex ratios were similar across treatment combinations. Because of the potential for birds to remain infected from prior exposure treatments until the time of secondary challenge (42 days post-prior exposure), we included a set of controls (n=10 total birds) that received prior exposure doses of pathogen (either low or high-dose) and then received a control inoculation of sterile media at day 42. These 10 birds were all qPCR negative for MG infection by day 41 and during all three subsequent sample periods post-secondary challenge (see Supplement).

Because our main interest was in how prior exposure treatment alters host susceptibility (1|0) to secondary dose challenge, we generally do not analyze or present data collected prior to secondary challenge (day 0), with one exception. To improve our dose response model fits for the “no prior exposure” treatment group, we included susceptibility data generated in response to the initial, low-dose prior exposure treatment. We use only the low-dose prior exposure treatment for this purpose, and not the high-dose prior exposure treatment (30000 CCU / mL) because the low-dose treatment uniquely provided an exposure dose (750 CCU / mL) that fell intermediate to those used in our secondary challenge doses (which were otherwise 30, 100, 300, and 7000 CCU/mL), informing a key part of the dose response curve. Thus, each of the 50 finches given a low-dose prior exposure treatment and then one of five secondary challenge doses (Table S1) contributed two data points to the analyses: (i) their susceptibility (1|0) to the “no-prior exposure” dose of 750 CCU/mL, with data collected on days −42 to −28 prior to secondary challenge; and (ii) their susceptibility (1|0) to one of five secondary doses, with data collected on days 4-14 post-secondary challenge. All other birds contributed only one data point to the data-set: susceptibility (1|0) to one of five secondary doses, with data collected on days 4-14 post-secondary challenge.

### Final sample sizes

We began with a balanced design (n=12 birds for all non-0 MG challenge doses in Table S1; n=157 total birds), but one individual died of unknown causes prior to secondary dose challenge. A second individual was excluded from all analyses after study completion because, when ELISA assays on frozen blood samples were completed, that individual was discovered to have been MG-seropositive just before prior exposure treatments were given. Finally, five additional individuals were later determined to be qPCR positive one day prior to secondary challenge (see *Susceptibility and disease sampling*), and thus were eliminated from all analyses to ensure that our susceptibility data represent true responses to secondary challenge rather than residual effects of prior exposure treatment. Final sample sizes for all analyses were n=150 individuals (Table S1).

### Susceptibility and disease sampling

We quantified whether birds were susceptible to infection or not (Y/N, 1|0) at a given MG exposure dose via qPCR of conjunctival swab samples collected post-exposure. We also visually scored clinical signs of conjunctivitis (all scoring was done blind to a bird’s treatment) to use as a proxy for the likelihood of mortality during infection (see below). At each sampling point, we first scored clinical signs of conjunctivitis on a scale of 0 to 3 for left and right conjunctiva separately, as per (86), with 0.5 score intervals used when clinical signs were intermediate between two integer scores. We summed the scores (left plus right conjunctiva) within a given sample day for a total maximum score of 6 for a given bird per day. After scoring, each bird’s conjunctiva was swabbed for 5s with a sterile cotton swab dipped in tryptose phosphate broth (TPB). Swabs were swirled in 300uL of sterile TPB and then wrung out into the sample collection tube. Samples from both eyes were pooled within sampling date for a given individual and frozen at −20°C until processing. Genomic DNA was extracted from samples using Qiagen DNeasy 96 Blood and Tissue kits (Qiagen, Valencia, CA) as previously described (84). Extracted genomic DNA from each sample was used to measure overall numbers of MG present in the conjunctiva using a qPCR assay targeting the *mgc2* gene of MG using the primers from (87) and qPCR methods outlined in (84).

Conjunctival swab samples and clinical scores were taken on days 4, 7, 14, and 21 post­secondary-challenge (and on days 7 and 14 post-prior exposure treatments). However, we only use conjunctival swab data from the first three sample points (days 4, 7, or 14 post-challenge) as most relevant for quantifying susceptibility. To account for potential low-level contamination in our qPCR assay (see Supplement) and the possibility that MG loads from prior exposure treatment might still be present in some birds at the time of secondary dose challenge, we considered a bird as susceptible to a given dose if they had a conjunctival MG pathogen load > 50 copies at any of the sample points from day 4 to 14 post-challenge. This cut-off fell above the highest MG load detected among birds that were given a sham secondary challenge treatment (but had been previously exposed), and thus should not encompass low MG loads that represent residual loads from prior exposure treatments. We also sampled all birds in the study on day 41 post-prior exposure, which was one day prior to secondary challenge treatment (day 42). As noted above, we eliminated 5/155 birds deemed infected on day 41 from all analyses (see Supplement).

### Dose response models and parameterization

We used dose-response models to describe the susceptibility of naive and previously exposed birds challenged with MG (20,46). We fit the models to data by minimizing the deviance between the data and the model and compared models using likelihood ratio tests that accounted for differences in the number of parameters between models (46). We assumed heterogeneous susceptibility followed a gamma distribution (Equation 1) (10). We compared models of homogeneous susceptibility (best supported for the no prior exposure group), with models of gamma distributed susceptibility (best supported for the two prior exposure groups). With these models we determined the proportion of individuals infected, *I_!om_* or *I_het_* via

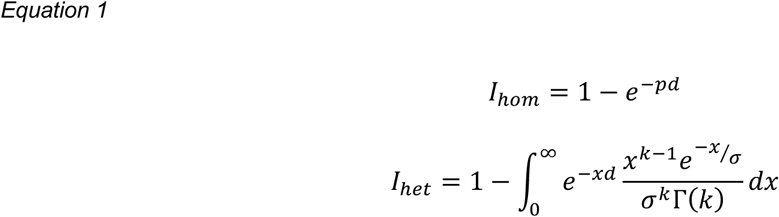

Here, *d* (dose) is the exposure concentration in CCU//Æ In *I_hom_*, the subscript *hom* denotes a model assuming homogeneous susceptibility with *p* representing the host infection probability per 1000 bacterial particles.The model with subscript *het* assumes individuals have susceptibility (x) that is distributed according to a gamma distribution with shape parameter *k* and scale parameter σ. Here, r(k) is the gamma function evaluated at the shape parameter k. We calculated coefficients of variation for the gamma models as the standard deviation divided by the mean.

### Fitting mortality data

We use clinical scores of conjunctivitis collected post-secondary challenge (with scores per bird per day ranging from 0 to 6; see above) to estimate mean mortality rates in the wild for each prior exposure treatment group. We assume that these ordinal scores monotonically predict host mortality because several lines of evidence indicate that increasingly severe conjunctivitis and resulting visual and behavioral impairment result in higher mortality risk for house finches in the wild: 1) the presence of conjunctivitis is linked with higher mortality in wild birds (52); 2) higher conjunctivitis severity predicts faster capture time in mock-predation events (i.e., a human capturing the bird by hand) in captivity (88); and 3) indirect mortality from predators appears to be the primary source of mortality for infected birds in the wild, because birds in captivity (a predator-free environment) do not ever succumb to infection, even with highly virulent strains (84). Thus, MG infection *per se* does not directly cause mortality in house finches, but mortality in the wild (via predation or the inability to find food) is directly linked with visual impairment and the associated disease that we measure.

Mortality rate was fit using nonlinear least squares regression (nls function in R) as the nonlinear inverse logit function, 1/(1+exp(-z)) scaled by 6, where z is the eye lesion score (integers between 0 to 6). This fit was then scaled by vmax (0.25; maximum mortality rate observed in the field from (52)) such that the maximum mortality rate is vmax rather than 1. A similar approach was used in previous work (34).

### Epidemic model

We use a Susceptible-Infectious-Recovered (SIR) model to examine the effect of heterogeneity in susceptibility on outbreak size and impact. For populations with determined heterogeneity in susceptibility, we consider a continuous distribution of susceptibility, with susceptibility, *x*, distributed according to the parameters (shape *k* and scale o) of the gamma distribution obtained from the dose response models. We model this with the differential equation system

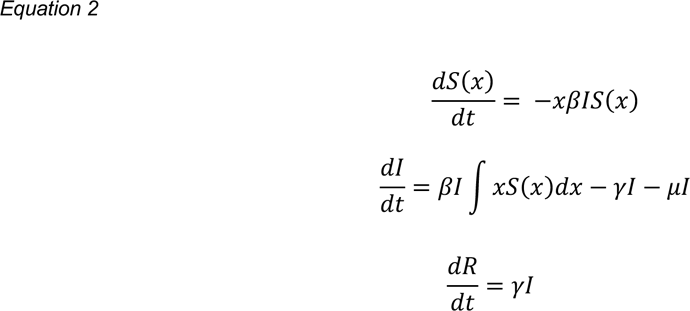

Here, *S(x)* is the number of susceptible individuals with susceptibility *x,* and *x* is a value between 0 and infinity (note that for simulation purposes, we truncate the integral at 10, which covers more than 95% of the full susceptibility distribution with our various choices of parameters). *I* is the number of infectious individuals, and *R* is the number of recovered individuals. Parameters for infectivity (/?) and recovery (γ) were taken from a prior empirically-parameterized model in this system (89) (Table S2), while disease-induced mortality (/) was fit as described above (See *Fitting mortality data*) for each prior exposure group (none, low, high). Infectivity and recovery rates do not depend on prior exposure, while mortality rate does (in the “fitted mortality” model). A density-dependent rather than frequency-dependent transmission term was used due to the nature of transmission in this system, which increases with density (76,90).

For simulation, we discretize the susceptibility distribution of the heterogeneous model into 300 evenly spaced susceptibility classes, represented by the midpoint susceptibility, x, for each class. We assume a population size of 100 and seed infections with a single infected individual. The initial population in each susceptibility class is determined empirically from the fitted distributions by prior exposure treatment, discretized into the 300 evenly spaced susceptibility classes which are truncated above susceptibility of 10 (See *Dose response models and parameterization* for details on determination of the fitted distributions). We quantify the total size of the epidemic as the cumulative fraction of infected individuals, and simulate for 500 days, after which nearly all epidemics have completely resolved. Because we simulate a short­time scale single epidemic, we ignore demographic effects and assume recovered individuals cannot be reinfected, despite documented waning of immunity to MG over the course of a year (53).

To disentangle the effects of heterogeneity in susceptibility and changes in mean susceptibility, we simulate outbreaks using the homogeneous model with identical mean susceptibility to the discretized heterogeneous distribution. To generate confidence intervals on outbreak sizes, we used the parameter estimates obtained from bootstrapping the chi-squared residuals of the dose-response estimates. We then estimated the probability of obtaining an epidemic size difference as extreme as observed (hom-het) using the best fit parameters, but with the heterogeneous case resulting in larger epidemics (het-hom).

## Data Availability Statement

The code and datasets generated for this study are available in the GitHub repository, at https://github.com/klangwig/mg_dose_response_public.

## Acknowledgments

We thank Edan Tulman for input on experimental design and MG isolate preparation, Chava Weitzman, John Brule, and many other Hawley lab members for help with capture and sampling, and Bambi Kelly for assistance with animal care. Research reported in this publication was supported by the National Institute Of General Medical Sciences of the National Institutes of Health under Award Number R01GM144972. The content is solely the responsibility of the authors and does not necessarily represent the official views of the National Institutes of Health.

## Author Contributions

D.M.H.: Conceptualization, Methodology, Resources, Investigation, Writing- Original Draft, Supervision, Project administration, Funding acquisition

A.P.U.: Investigation, Data Curation, Writing - Review & Editing

J.S.A: Conceptualization, Methodology, Writing-Review and Editing, Funding acquisition

AEFD: Conceptualization, Methodology, Software, Formal Analysis, Writing - Review & Editing, Funding acquisition

J.N.GL: Investigation, Writing-Review and Editing

S.J.G.: Conceptualization, Methodology, Writing-Review and Editing

L.M.C.: Conceptualization, Methodology, Software, Validation, Formal Analysis, Writing-Review and Editing, Visualization, Funding acquisition

K.E.L.: Conceptualization, Methodology, Software, Validation, Formal Analysis, Writing-Review and Editing, Visualization, Funding acquisition

