## Supplemental Materials for "Prior exposure to pathogens augments host heterogeneity in susceptibility and has key epidemiological consequences"

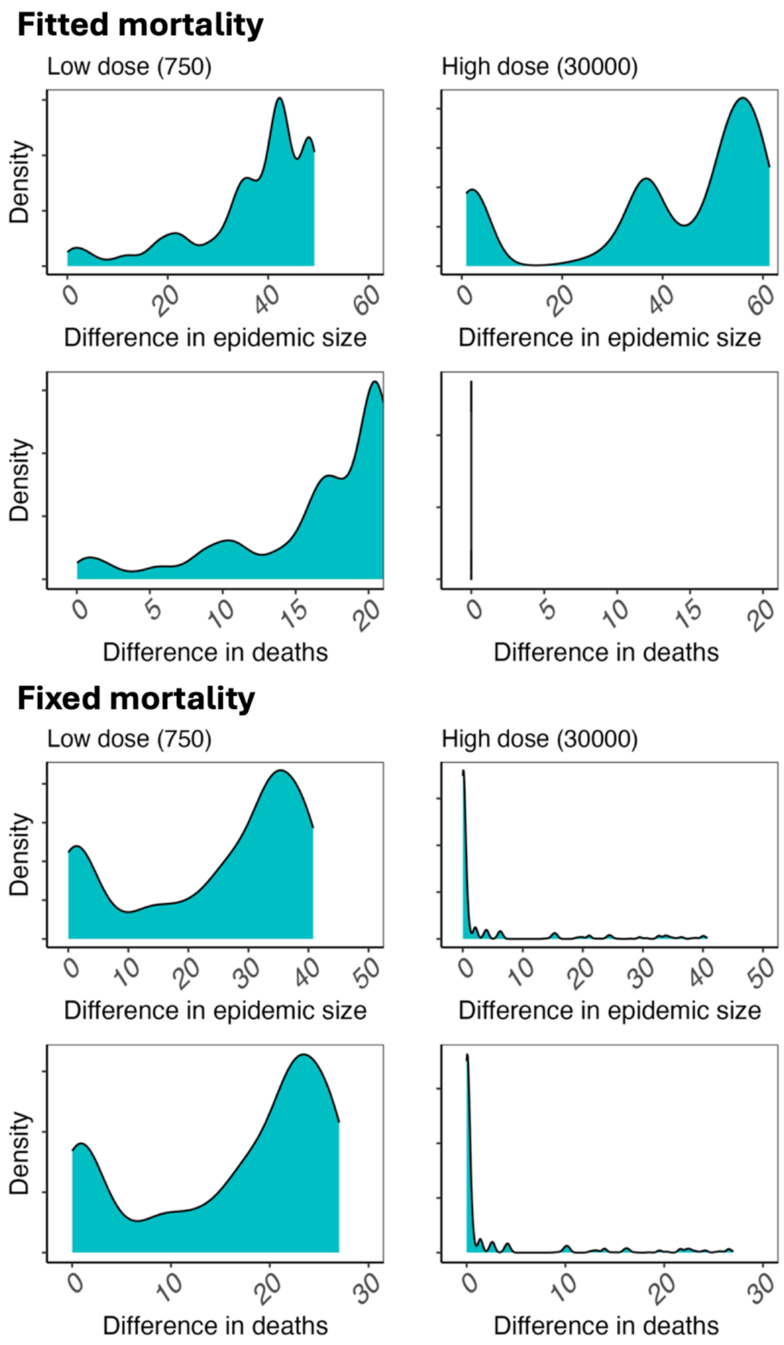


**Figure S1.** Density plots of observed increases (difference=homogeneous - heterogeneous value) in both epidemic size (left) and total mortality (right) for simulations using parameter estimates obtained from bootstrapping the chi-squared residuals of the dose-response parameters (Fig 3). Paired comparisons were made while holding mean susceptibility constant, and only varying whether susceptibility distributions were homogeneous versus heterogeneous (fitted parameters from each bootstrap sample). The top four panels are from simulations using mortality rate parameters fitted to empirical data for each prior exposure treatment; bottom four panels are from simulations using a fixed mortality rate (that of the no prior exposure group)

across all prior exposure treatments. Density plots are scaled to the maximum height across a fixed x-axis scale (for fitted mortality, maximum of 60 for epidemic size and maximum of 20 for deaths; for fixed mortality, maximum of 50 for epidemic size and maximum of 30 for deaths). Confidence intervals for epidemic size differences are reported in the main text (see *Results*). For differences in deaths (hom-het), 95% CIs were: low dose, fitted mortality [0.9906,23.74]; high dose, fitted mortality [0,0]; low dose, fixed mortality [0.07863,27.01]; and high dose, fixed mortality [0.0064,24.09].

**Table S1.** Experimental design testing how prior exposure to *M. gallisepticum* (none, low dose, high dose) alters host heterogeneity in MG susceptibility (n=157 at start; n = 150 for final analysis). We began with a balanced design for all non-0 secondary MG doses, but lost one bird to mortality and excluded six birds retroactively, one of which was discovered to be seropositive to MG at start, and five that were discovered to be qPCR positive one day prior to secondary challenge (see Methods).

| **Prior Exposure Treatments** | **Secondary MG Dose Treatments (CCU/mL)** | | | | |
|---|---|---|---|---|---|
| | **0** | **30** | **100** | **300** | **7000** |
| Sterile media (No prior MG exposure) | n=3 | n=12 | n=12 | n=11 | n=12 |
| Prior low dose MG (750 CCU/mL)* | n=4 | n=11 | n=12 | n=11 | n=12 |
| Prior high dose MG (30,000 CCU/mL) | n=6 | n=11 | n=12 | n=11 | n=10 |

*Whether birds became infected or not in response to the **prior low MG dose treatment** (n=50; gray cells) was used to estimate susceptibility of birds with no prior exposure, which was the case for these birds at the time of this treatment**.**

**Table S2.** Parameters used in our SIR model to examine effects of host prior exposure on epidemiological dynamics. For all heterogeneous SIR simulations, gamma-distributed susceptibility values (first row) were used to determine the initial susceptibility distribution, while for all homogeneous SIR simulations, susceptibility values (second row) were used to determine the susceptibility of the entire population. The susceptibility for homogeneous simulations was determined from the mean of the corresponding heterogeneous models. Transmissibility and recovery rates for this system (obtained from[1]) were held constant across all groups, while other parameters were varied using model fits to empirical data. *In simulations with fixed mortality rates across groups, we used the mortality rate estimated for the no prior exposure group.

| **Parameter** | **No prior exposure** | **Low-dose prior exposure** | **High-dose prior exposure** |
|---|---|---|---|
| Heterogeneous (gamma-distributed) susceptibility, x | shape = 1.237<br>scale = 0.955<br>(mean = 1.181) | shape = 0.376<br>scale = 1.185<br>(mean = 0.4460) | shape = 0.159<br>scale = 1.21<br>(mean = 0.1920) |
| Homogeneous (exponential) susceptibility, x | x = 1.181 | x = 0.4460 | x = 0.1920 |
| Fitted mortality rate | 0.05894* | 0.02806 | $2.483*10^{-17}$ |
| Transmissibility | 0.00275 | | |
| Recovery rate | 0.03 | | |

**Supplemental Methods**

**Sampling timeline**
We collected baseline samples (blood from the brachial vein, and conjunctival swabs) from all individuals 7-8 days before prior exposure treatments. We used these samples to ensure that birds were seronegative and pathogen negative just prior to experimental treatments. Blood samples were also taken one day prior to secondary challenge (data not presented), and conjunctival swab samples were taken on days 7, 21, and 41 post-prior exposure treatment. A single individual assigned to the sham treatment (that was also known to be seropositive; see above) was qPCR positive (169.90 pathogen copies) the week prior to the experiment beginning. Because this individual was known to be seropositive (and was not used in any MG-treatment groups), this positive value was assumed to represent residual infection from the wild. This individual was not detected as "infected" once the experiment began.

**Quantifying Susceptibility.** Our qPCR assay is highly sensitive, and because we use 96-well plates for extractions, low level cross-sample contamination cannot be completely eliminated in our assays (see ESM in[2]). We use three types of controls to account for such low-level contamination: 1) extraction controls, which are samples scattered throughout the 96-well extraction plate (3 to 5 per plate) that experience the entire extraction process but without any starting sample; 2) qPCR assay negative controls (three per assay), which experience the entire assay but without any starting template; and 3) swabs from sham control birds that we know were not experimentally exposed to MG in our experiment. Of the extraction controls analyzed for this study, two were positive (where positivity is defined by any signal detection of amplification in the qPCR assay) with very low estimated pathogen loads of <5 copies of MG. No qPCR assay negative controls were ever positive. Of the 21 extracted samples from the sham birds that had never been experimentally exposed to MG (but were seropositive at capture), two were positive, with estimated pathogen loads of 1.70 and 169.90 copies (the latter of which was detected prior to the experiment officially beginning, as noted above). To account for the known high specificity and potential background noise in our assay, we considered an individual as susceptible if they met a conservative threshold of ≥50 copies of MG present in the conjunctivae at any timepoint post-challenge (see main text).

We also included a set of controls (Table S1) given prior exposure to MG at low (n=4) or high-dose (n=6), but then given a control secondary challenge treatment (sham sterile media, represented as 0 MG in Table S1). These 10 controls were included to account for potential lack of recovery from prior exposure treatments before secondary challenge treatment. None of these 10 birds were deemed "infected" on day 41, just prior to secondary challenge (day 42). Further, none of the 10 birds were positive at our infection threshold (≥50 copies of MG ) at any sampling timepoint following secondary challenge (on days 4, 7, or 14 post-secondary treatment). This suggests full clearance of MG from prior exposures by the time of secondary dose challenge assay for all birds in this control group. Nonetheless, outside of this control group, five finches (out of 155 total in the study) were deemed "infected" via our 50-copy cut-off at day 41, which was one day prior to secondary dose challenge (day 42). Thus, to be as conservative as possible in defining susceptibility to secondary challenge, all five of these birds were eliminated from all analyses (Table S1), for a final sample size of n=150 for analysis.

**Housing**
All birds were fed an *ad libitum* diet of Roudybush Daily Maintenance Pellets Nibles with sunflower kernels mixed in at approximately 20% of the pellets. *Ad lib* drinking water was provided at all times. After a minimum two-week quarantine and at least 10 days before prior exposure treatments, all birds were housed individually in wire-mesh cages (18"x18"x30") in one

of seven indoor rooms that varied in size but experienced identical daylengths (12:12 light:dark) and temperature ranges. To control for potential variation among housing rooms related to size, we ensured that rooms contained approximately equal proportions of each treatment group and sex ratios. On 29 Oct 2021 (day 10 post-prior exposure treatment), all birds in one of the seven housing rooms had to be moved unexpectedly to a new nearby housing room (but remained in their home cages, with the move taking approximately 10 minutes in length) due to a sudden temperature spike in their original housing room that required repairs. To minimize further disturbance, birds remained in the new housing room for the remainder of the study. Because all treatments and sexes were evenly distributed among housing rooms at the start of the study, this move should not have any effect on our results or interpretation.

For the "sham" control birds that were not exposed to *Mycoplasma gallisepticum* (MG) at either experimental timepoint (n=3, Table S1), we used birds that were seropositive via ELISA on days 14-18 post-capture (see Methods in main text). Because these three sham control birds only served as sham controls for handling procedures, the use of seropositive birds as controls has no effect on our conclusions and allowed us to minimize the overall collection and use of wild-caught birds. Two of the three "sham" control birds were housed in a pair for the prior exposure portion of the study only, and then moved to single cages thereafter. All other birds were housed individually throughout the duration of the study.

**Prophylactic medications**

Beginning in the first two weeks following capture, all birds were treated with several prophylactic medications (all dosed in the drinking water) to prevent mortality from known pathogenic agents that affect captive finches. First, within 10 days post-capture, all birds were treated with Cankerex Plus (Dimetridazole B,P (Vet) 40% m/m powder) dosed at 0.25g/L for 5 days in the water to prevent trichomoniasis. Following completion of Cankerex treatment, all birds were treated with Endocox (2.5% Toltrazuril) at a dose of 1.32 g/ L in the water to help control coccidia overgrowth. Endocox treatments were given in 3-day pulses every week (3 days on/4 days off) for 4 consecutive weeks, followed by monthly or bimonthly 3-day pulses thereafter. On 3-4 days per week that birds did not receive Endocox, birds were given water with probiotics at a concentration of 1 g/ L (Bene-Bac Plus Bird and Reptile Supplement; Item No. 186877).

In response to unexpected mortality in mid-July 2021 that was a suspected result of gastrointestinal fungal overgrowth, we began adding a low concentration of store-bought apple cider vinegar (5 mL / L) to the water of all birds 4 days / week (approximately every other day). Apple cider vinegar additions were ceased on 1 October 2021, over 2 weeks prior to the experiment beginning. Due to a second round of unexpected mortality in late August 2021, birds in two quarantine housing rooms (out of 7 total rooms) were given a round of Nyastatin treatment in their feed (5g nystatin powder/1 kg seed) for 7 days (beginning 31 Aug 2021) to prevent GI fungal overgrowth. Because such treatment did not appear to reduce observed unidentified mortality (which is known to occur regularly when hatch-year wild birds are brought into captivity), other housing rooms were not treated. Although Nyastatin treatment ceased 6 weeks prior to initiation of the experiment, to be conservative, we ensured that birds that received Nyastatin treatment were randomly assigned across all treatment groups, such that each treatment group had roughly equal proportions of birds that did or did not receive Nyastatin prior to the experiment. All other prophylactic medications were identical across individuals throughout the course of quarantine and the experiment.
